## Supporting Information for "The Role of Adaptation in Generating Monotonic Rate Codes in Auditory Cortex"

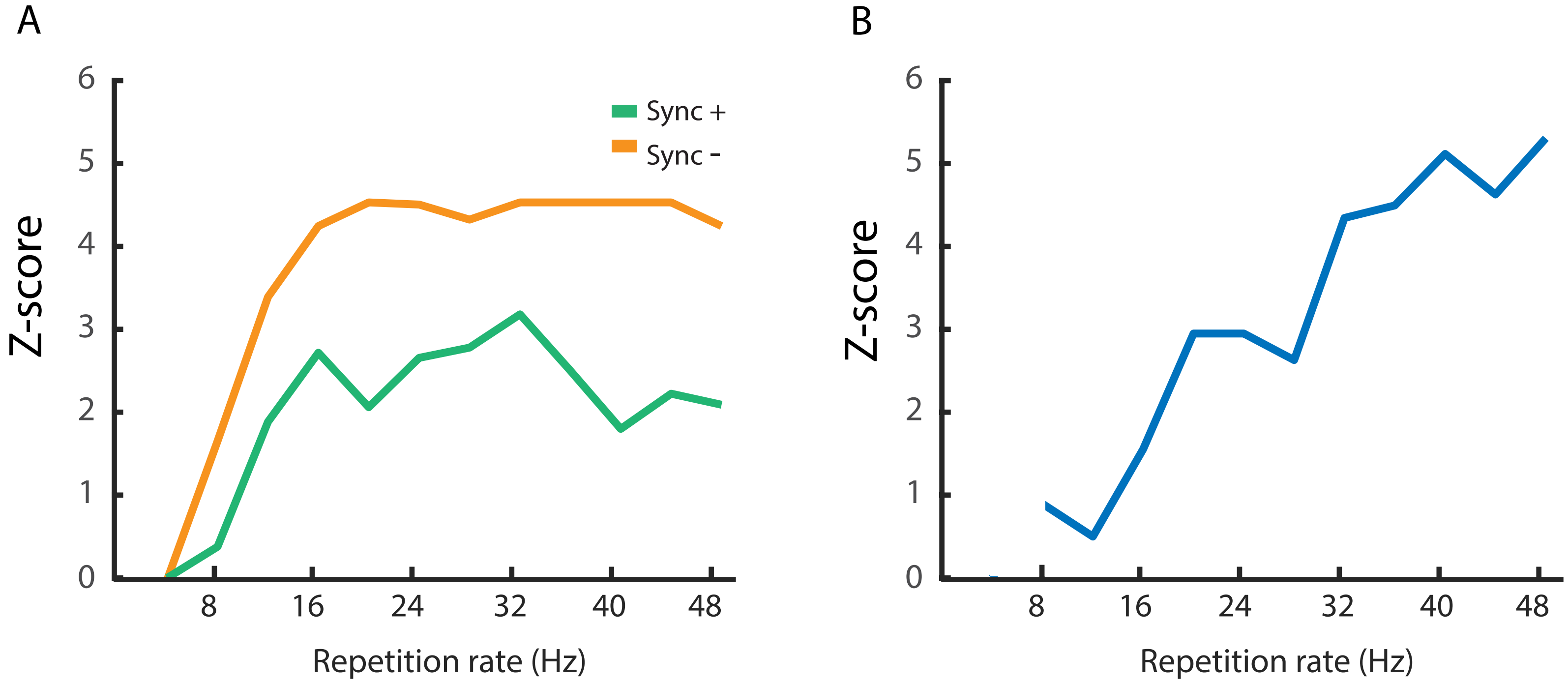


**S1 Fig.** Adaptation to stimuli in Sync+ and Sync- neurons.

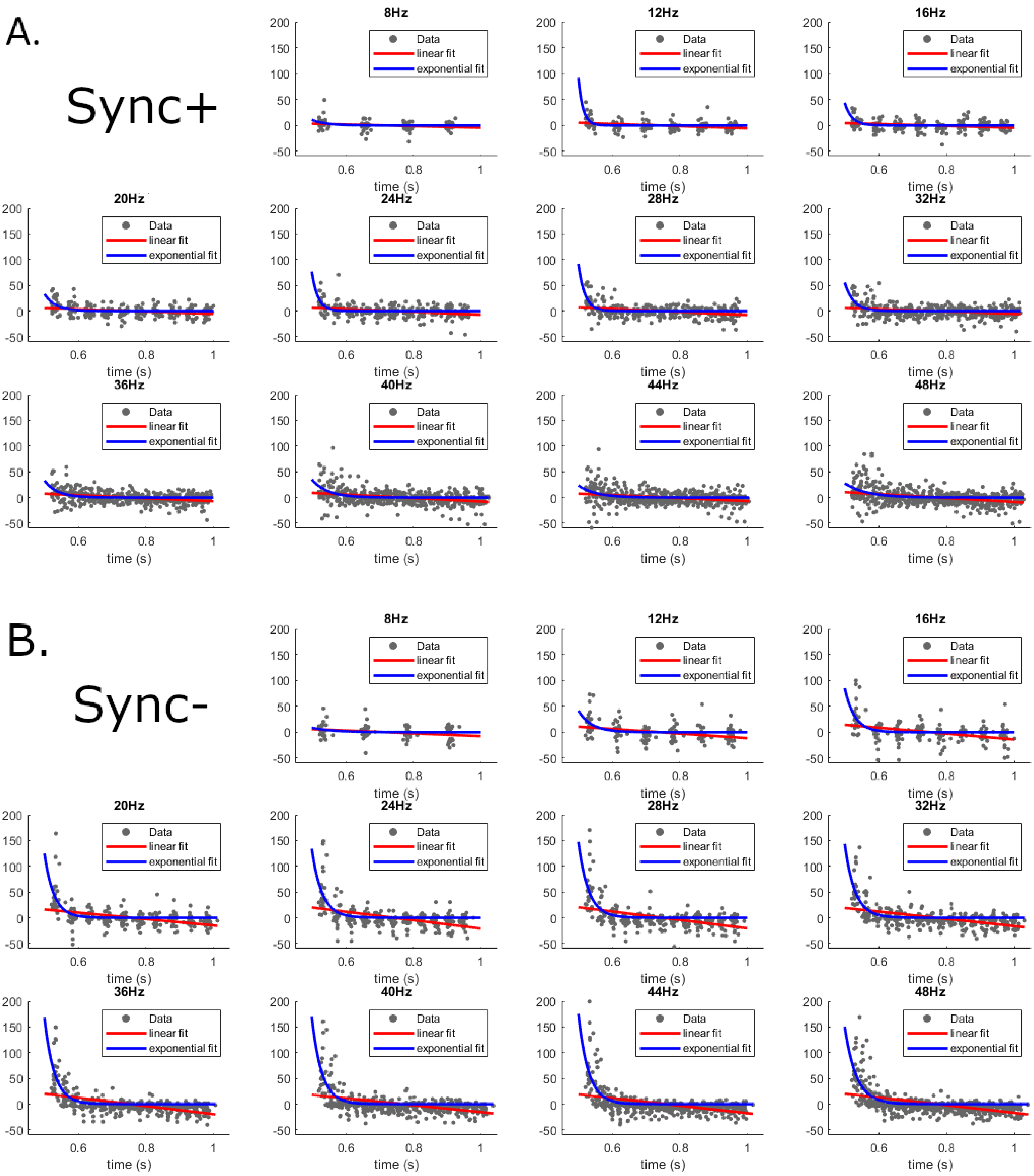


**S2. Fig.** Sync+ (a.) and Sync- (b.) neuron responses to stimulus pulse trains

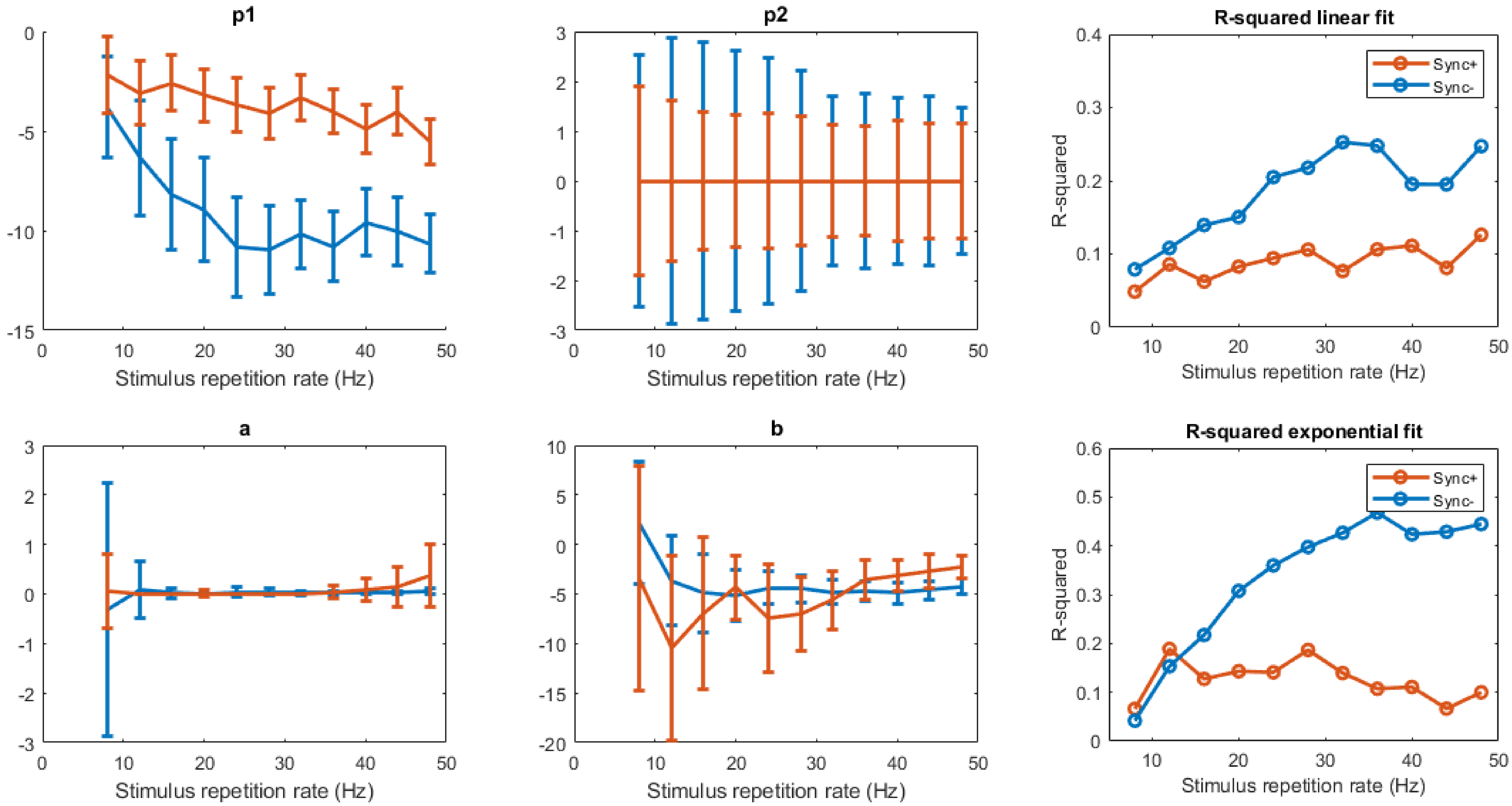


**S3 Fig.** Fitted model coefficients to adaptation during stimulus presentation

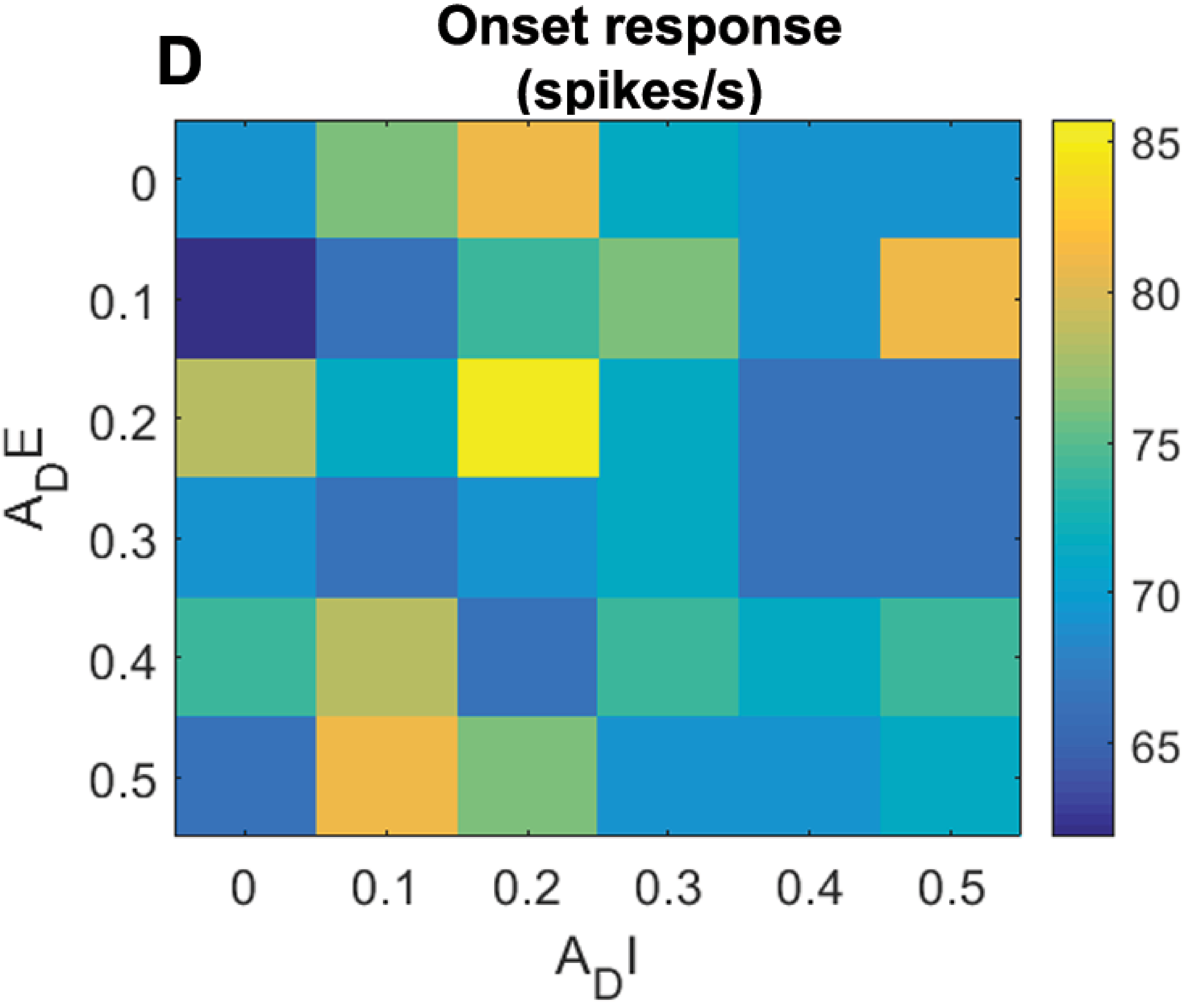


**S4 Fig.** Onset response amplitude relative to strength of adaptation.

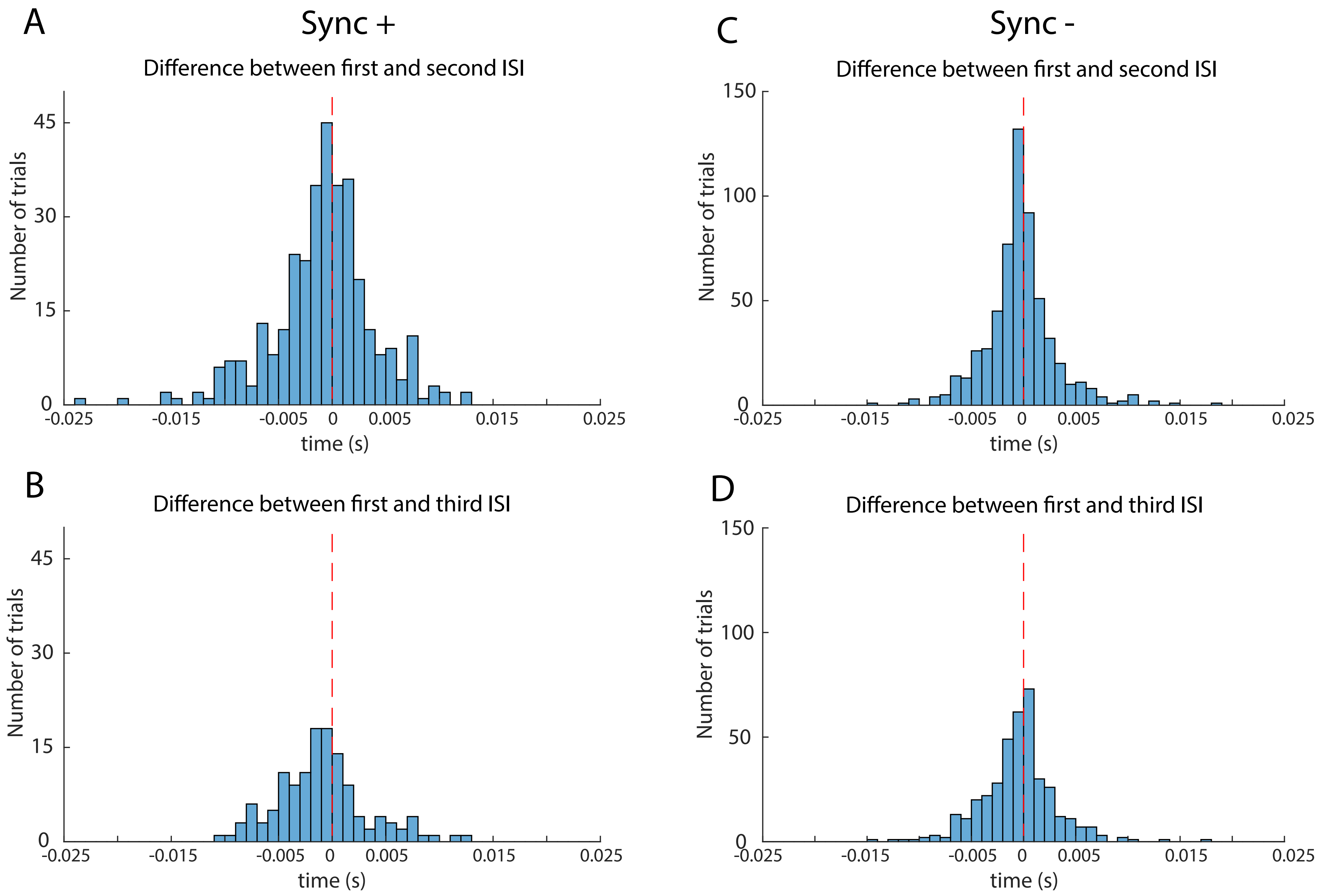


**S5 Fig.** Comparison of ISI after stimulus onset

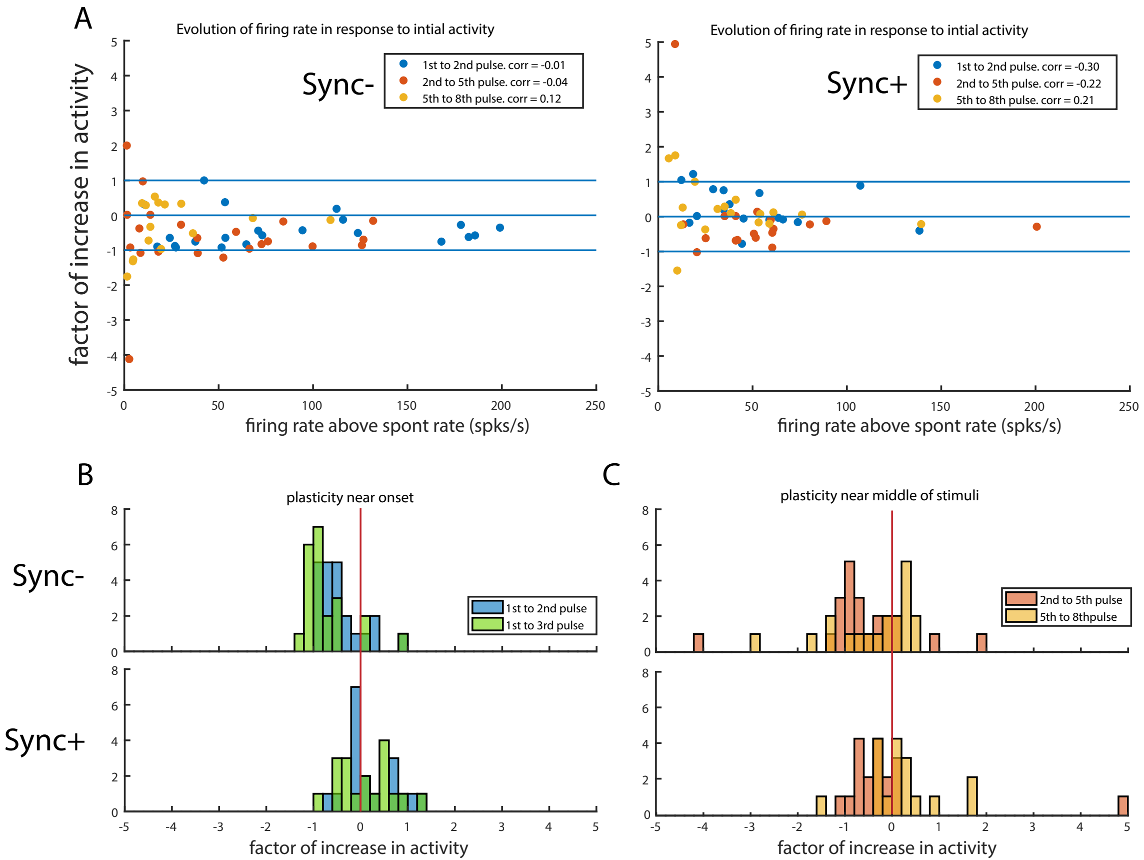


**S6 Fig.** Monotonicity and adaptation in individual neurons

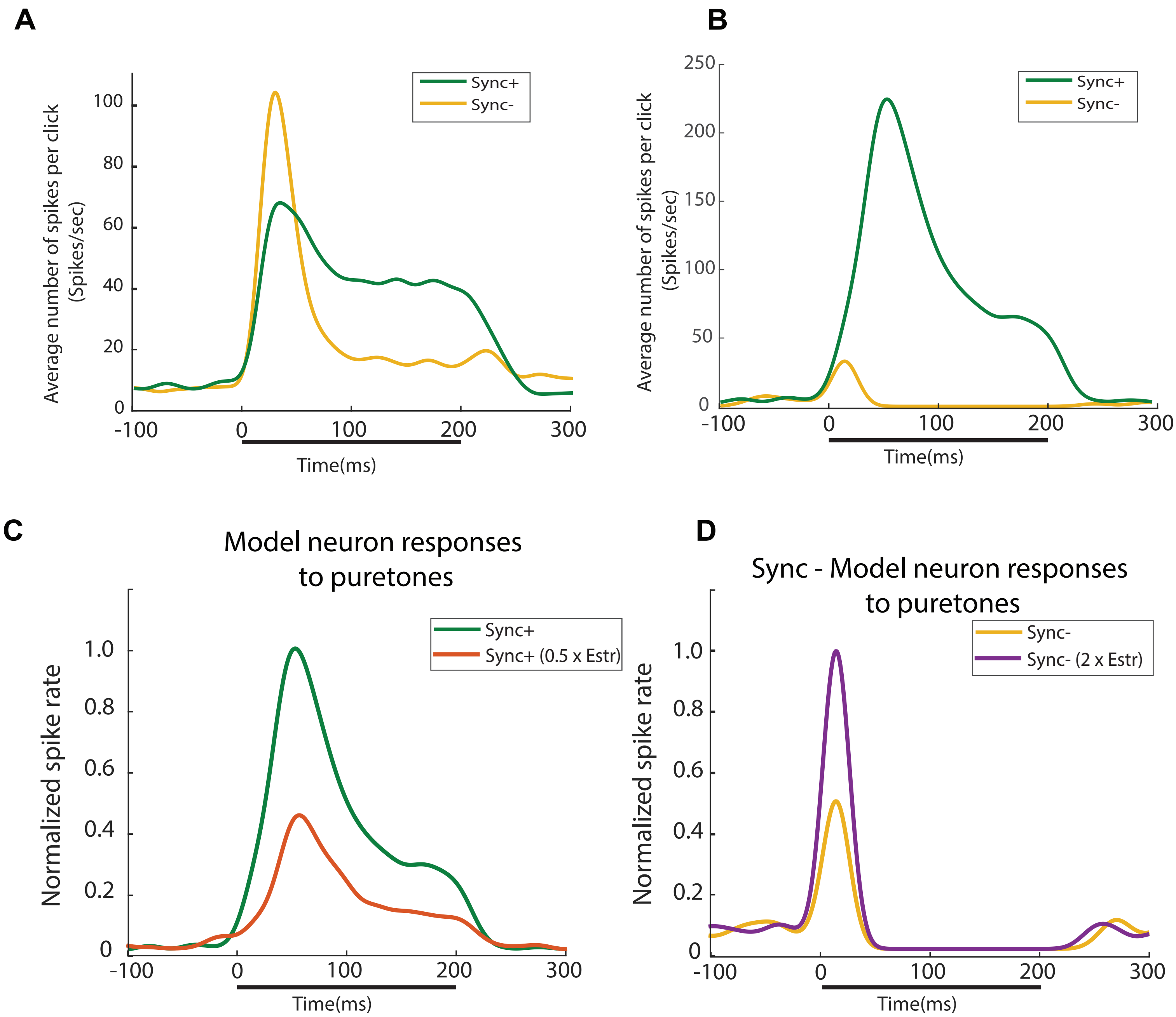


**S7 Fig.** Puretone responses

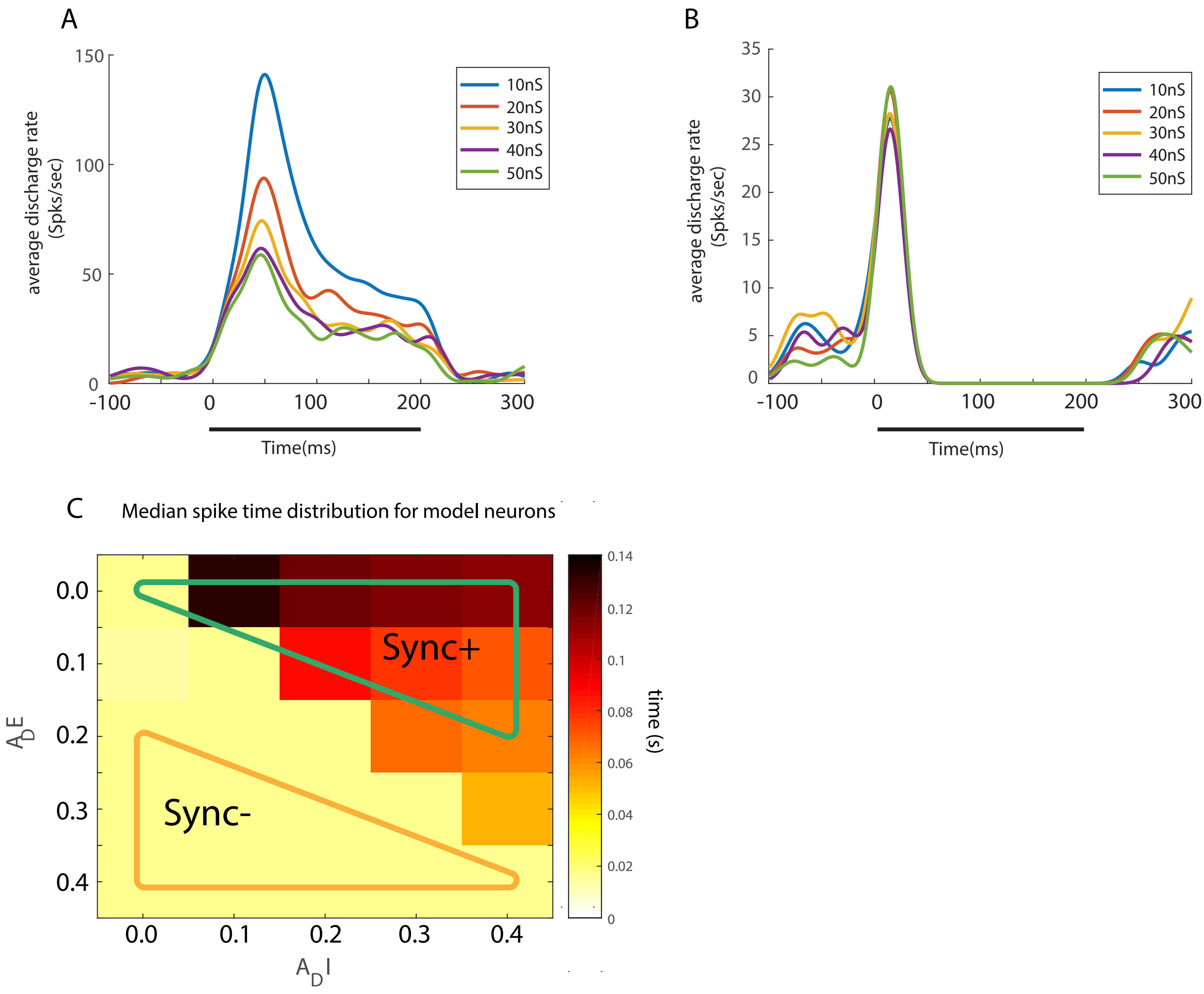


**S8 Fig.** Puretone responses and SFA.
